# Reversion is most likely under high mutation supply, when compensatory mutations don’t fully restore fitness costs

**DOI:** 10.1101/2020.12.28.424568

**Authors:** Pleuni S. Pennings, C. Brandon Ogbunugafor, Ruth Hershberg

**Affiliations:** Department of Biology – San Francisco State University, San Francisco, CA, 94132 USA; Department of Ecology and Evolutionary Biology – Yale University, New Haven, CT, 06520 USA; Rachel & Menachem Mendelovitch Evolutionary Processes of Mutation & Natural Selection Research Laboratory, Department of Genetics and Developmental Biology, the Ruth and Bruce Rappaport Faculty of Medicine, – Technion-Israel Institute of Technology, Haifa, 31096 Israel

**Author notes:** The authors contributed equally to this work.

## Abstract

Adaptive mutations are often associated with a fitness cost. These costs can be compensated for through the acquisition of additional mutations, or the adaptations can be lost through reversion in settings where they are no longer favored. While the dynamics of adaptation, reversion and compensation have been central features in several studies of microbial evolution, few studies have attempted to resolve the population genetics underlying how and when either compensation or reversion occur. Specifically, questions remain regarding how certain actors—the evolution of mutators and whether compensatory mutations alleviate costs fully or partially— may influence evolutionary dynamics of compensation and reversion. In this study, we attempt to explain findings from an experimental evolution study by utilizing computational and theoretical approaches towards a more refined understanding of how mutation rate and the fitness effects of compensatory mutations influence evolutionary dynamics. We find that high mutation rates increase the probability of reversion towards the wild type when compensation is only partial. However, the existence of even a single fully compensatory mutation is associated with a dramatically decreased probability of reversion to the wild type. These findings help to explain specific findings from experimental evolution, where compensation was observed in non-mutator strains, but reversion (sometimes with compensation) was observed in mutator strains, indicating that real-world compensatory mutations are often unable to fully alleviate the costs associated with resistance. Our findings emphasize the potential role of the supply and quality of mutations in crafting the evolution of antibiotic resistance, and more generally highlight the importance of population genetic context for explaining findings from experimental evolution.

## INTRODUCTION

In experimental evolution with microbes, rapid adaptation to new environments is often achieved through mutations displaying antagonistic pleiotropy [1]–[6]. This means that while such mutations are adaptive for specific selected traits, the fitness improvement often comes at the expense of other traits [3], [4], [7]–[10]. For example, a population of bacteria that is resistant to antibiotics may be less fit (compared to the susceptible ancestor) in an environment without antibiotics. Therefore, when the drug is removed, the mutations conferring resistance are no longer beneficial and may incur a fitness cost. One naive expectation is that when antibiotic usage ceases, the population will evolve “backwards” towards being drug susceptible. However, theory tracing back to paleontology (Dollo’s Law) has speculated that reversal to ancestral genotypes can be infrequent and challenging to achieve [11]–[13].

The question of reversal is relevant to the biomedical paradigm, where attempts to undo resistance by halting the use of antimicrobial drugs have been effective in some contexts [14], [15], but unsuccessful in others [7], [16]. The mixed success suggests that the plausibility of reversion is contingent on certain population genetic specifics.

Many studies have examined the persistence or reversion of costly adaptations by studying bacterial populations fixed for mutations conferring resistance to an antibiotic or phage (e.g. [7]–[9], [17]–[19]). These studies demonstrate that bacteria have a remarkable capability to compensate for the deleterious effects of resistance mutations by acquiring compensatory mutations, facilitating the maintenance of resistance [7]–[9], [17]–[19]. Further, such studies provide a mechanistic basis for the curious infrequency of empirical examples of reversion. For example, some studies have highlighted the role of non-linear forces like epistasis in undermining reversal across a fitness landscape [20], [21].

Though resistance to environmental stressors like antibiotic resistance and phage infection are great models, questions about the dynamics of reversion and compensation transcend bacterial resistance, and are of theoretical relevance within both microbial and non-microbial contexts. For example, recent studies have identified that the fitness effects of compensatory mutations are often partial in experimental evolution for cellular modules [22].

The model bacterium *Escherichia coli* can survive for very long periods of time within spent media (with no supplementation of external nutrients) in a state termed long term stationary phase (LTSP). Previous studies have demonstrated that *E. coli* adapts under LTSP through the acquisition of mutations within the RNA polymerase core enzyme (RNAPC) [3]. While these mutations are adaptive under LTSP they carry a cost that manifests in poorer growth in fresh media. Intriguingly, a recent study found that the tendency of RNAPC adaptations to persist within fresh media was related to the mutation rate [23]: non-mutator clones recovered through compensatory mutations, while mutator clones (∼ 110 − 260 fold higher mutation rates than the wild type, *WT*) that were able to recover near ancestral growth rates did so through reversion. Though these specific results were not rigorously explained, the general findings are in line with a literature supporting the role of mutation rate in dictating the dynamics of adaptive evolution in microbes [24]–[27].

In this study, we use theoretical and computational approaches to dissect the population genetic particulars underlying how variation in mutation rates affects the persistence of costly adaptations and the dynamics of compensation and reversion. In agreement with the long term stationary phase experimental results [23], we demonstrate that the likelihood for reversion in lieu of the persistence of resistance through compensation increases with higher mutation rates. Yet, even at high mutation rates, reversion is unlikely if there exist compensatory mutations that fully alleviate the costs associated with the adaptations (perfect compensation).

## MODEL AND METHODS

### Note on terminology: “reversal” and “reversion”

In this study, we use “reverse” and “reversion” interchangeably, and interpret them as the processes through which an allele reverts back to its ancestral wild type form.

### Note on simulations

Our simulations follow the population after it has adapted to a new (temporary) environment through the fixation of an adaptive mutation. Calculations and simulations start when the environment changes back to the original environment, and the adaptive mutation is now associated with a cost (c). For the remainder of the study, we will refer to the adaptive mutation as a resistance mutation. However, it is important to note that our results are relevant to any adaptive mutation that carries a cost.

Several types of mutations can occur in our model populations.

1. A reversion mutation removes the resistance mutation and recreates the original *WT* genotype. This mutation occurs with a probability of *µ* per generation and per bacterial cell. Bacteria with a reverted genotype are indicated by “*WT* ”.
2. Compensatory mutations reduce the cost of the resistance mutation by p such that the cost becomes *c*\* (1 − *p*). p is set to 0.5 in most of the simulations. We allow n different compensatory mutations to happen, where *n* is set to 100 for most simulations. Each of the n compensatory mutations occurs with a probability of *µ* per generation and per bacterial cell. Bacteria with a compensatory mutation are indicated by “Com” in the figures. In our simulations, we do not allow for bacteria to acquire multiple compensatory mutations.
3. Bacteria that carry a compensatory mutation can acquire a subsequent reversal mutation in addition to their compensatory mutation, with a probability of *µ* per generation and per bacterial cell (though this feature is turned off for some of the simulations). Bacteria that carry a compensatory mutation and are then reverted to *WT* are indicated by “compensated reversal” or “CR” in the figures.

### Additional notes on simulations and associated calculations

We carried out forward-in-time (Wright-Fisher) computer simulations using a custom R script. In the simulations, a population of size N (10, 000) individual bacteria is followed over G (100 or 500) generations. Each new generation is generated by multinomial sampling to include differences in fitness and the stochastic nature of reproduction. We also used a simplified deterministic approximation to calculate the probability of different outcomes (see supplementary material). All code is available at: https://github.com/pleunipennings/CompensatoryEvolution.

### Possible evolutionary outcomes

We created three illustrative scenarios, reflecting possible simulation runs (see Figure 1). Under all three scenarios, initially, a resistant allele (orange) rises in frequency due to strong selection until it is fixed in the population. This study focuses on what happens after the resistant allele is fixed, and when there is no longer selection for resistance. Will the population evolve back to wild type? Or will the cost or resistance be compensated instead?

**Figure 1:**
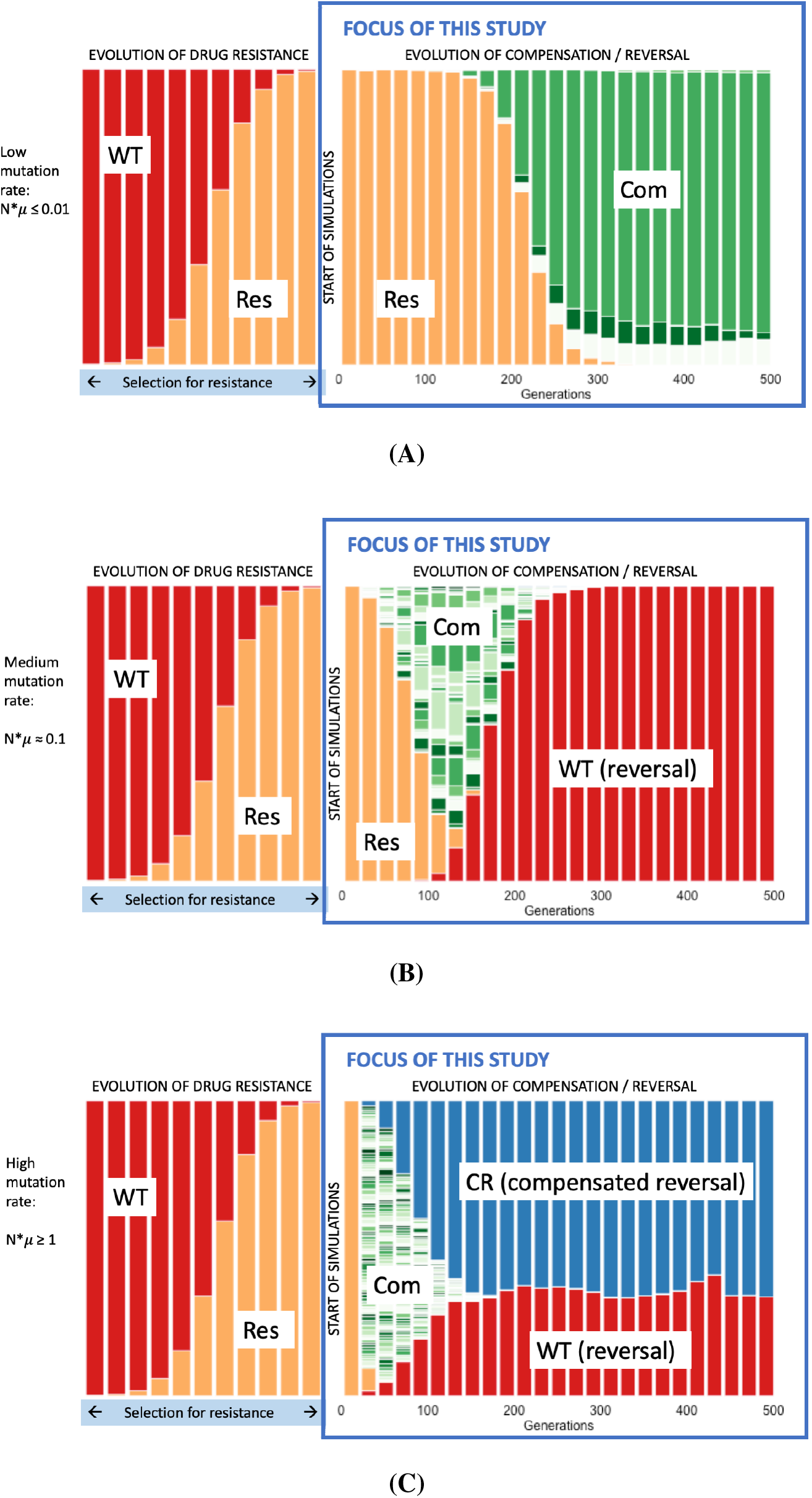
Illustrative simulations with different mutation supply rates. **(A)** Low mutation supply rate leads to compensation only in the first 500 generations. **(B)** intermediate mutation supply rate usually leads to *WT* outcompeting the compensatory mutations. **(C)** high mutation supply rate leads to *WT* reversion mutation occurring by itself (red) and on the background of a compensatory mutation (blue). The X-axis represents generations, and the Y-axis corresponds to the frequency of a given genotype in the population. Because there are multiple possible compensatory mutations, we have chosen to depict them with different shades of green. All compensatory mutations have the same compensatory effect. Red = *WT*, orange = resistant, blue = compensated and reverted.

In a first scenario (Figure 1A), the mutational supply in the population is low (one reversal mutation, at most, every 100 generations, *N*µ* ≤ 0.01). Compensatory mutations (of which there are *n*) mostly likely arise and fix in the population before a successful *WT* reversal mutation occurs.

In a second scenario (Figure 1B) there is an intermediate mutation supply in the population (approximately one *WT* reversal mutation every 10 generations, *N*µ* = 0.1). Compensatory mutations (green) likely arise in the first generation. However, none of the compensatory mutations can fully restore fitness, and in this higher mutation rate scenario, a *WT* reversal mutation (red) likely occurs before the compensatory mutations fix. The *WT* reversal mutation has higher fitness than the compensatory mutations and once established will fix in the population. The final scenario (Figure 1C) is one of high mutation supply rates (one or more *WT* reversal mutations every generation, *N*µ* ≥ 1). Here, a different pattern emerges: both compensation and *WT* reversal arise (as in Figure 1B), however, before the *WT* reversal mutations fix in the population, the bacteria with compensatory mutations also acquire *WT* reversal mutations that remove the resistance mutation. Now there are *WT* reversal mutations on two different genetic backgrounds: with and without a compensatory mutation. This scenario resembles a “soft sweep” where the reversal mutation occurs multiple times, both on a compensated genotype and on an uncompensated genotype [28]. Because these two *WT* reversal types have equal fitness in our model (and likely often in reality) compensated and uncompensated alleles can co-exist at intermediate frequency for a long time.

## RESULTS

### 1. Direct WT reversal is more likely when N*µ is higher, the mutational target for compensatory mutations is lower and when compensation is less effective

Firstly, we sought to determine the probability of the scenario outlined in Figure 1B (where a *WT* reversion mutation outcompetes compensatory mutations) vs the scenario in Figure 1A (compensatory mutations fix in the population). For simplicity, we do not allow *WT* reversion mutations to occur on an already compensated genotype (that is, scenario 1C is not possible).

*WT* reversion mutations have higher fitness than the compensatory mutations. Therefore, if a *WT* reversal mutation occurs and escapes genetic drift, it will outcompete the compensatory mutations. However, if the mutation supply rate in the population (*N*µ*) is low, it is possible and likely that the compensatory mutations fix before a successful *WT* reversal mutation occurs. We can see this situation as a race between fixation of the (very common) compensatory mutations and occurrence of a (rare) WT reversal mutation and there are only two possible long-term outcomes: either the entire bacterial population will consist of those with compensatory mutations, or with the reverted *WT* genotype.

We find that *WT* reversal mutations are most likely to occur before the compensatory mutations fix when the mutation supply is high (shorter wait time for a *WT* reversal mutation), when the mutational target size for compensatory mutations (*n*) is low or when the effect of compensation is small (*p*) (small values for these two parameters make the fixation of the compensatory mutations slower, which gives more opportunity for the *WT* reversal mutations to occur) (figure 2).

**Figure 2:**
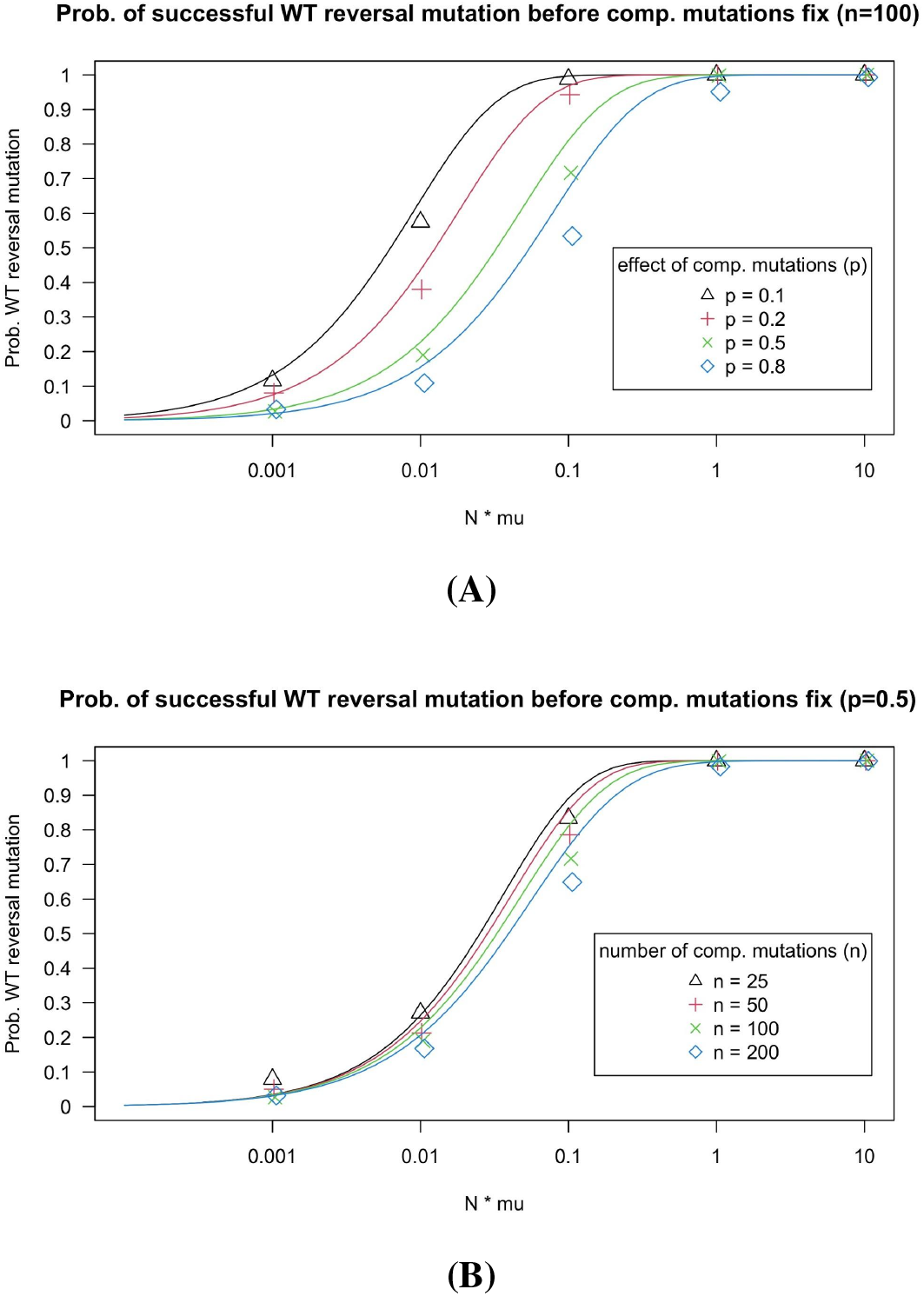
**(A)** The probability that a *WT* reversion mutation occurs before the compensatory mutations fix in the population. In these simulations, clones with compensatory mutations cannot acquire a second mutation (i.e., they cannot revert to *WT*). Simulations are run for different values of *µ*, the mutation rate and different values of p, the compensatory effect. Lines are based on the deterministic model. **(B)** Same as A, but simulations are run for different values of n, the target size for compensatory mutations. Simulations are run for 500 generations. Other parameters: *N* = 10, 000, *c* = 0.15.

Simulations agree well with a deterministic model (see supplementary materials and figure 2).

### 2. WT on background of compensatory mutation occurs when the mutation supply is high

When the mutation supply is high (*N*µ >* 1), the WT reversal mutation can occur on two different backgrounds. It can occur by itself (indicated by the red color in figure 1) or on a background that already carries a compensatory mutation (indicated by the blue color in figure 1). We show here that there is a surprisingly sharp transition between two regimes. If *N*µ* is 0.1 or lower, it is uncommon to see the WT reversal mutation on two backgrounds in the simulations. However, if *N*µ* is 1 or higher, we see it in almost all simulations (see figure 3). The effects of p (effect of compensatory mutations), n (number of compensatory mutations) and G (number of generations) are quite small.

**Figure 3:**
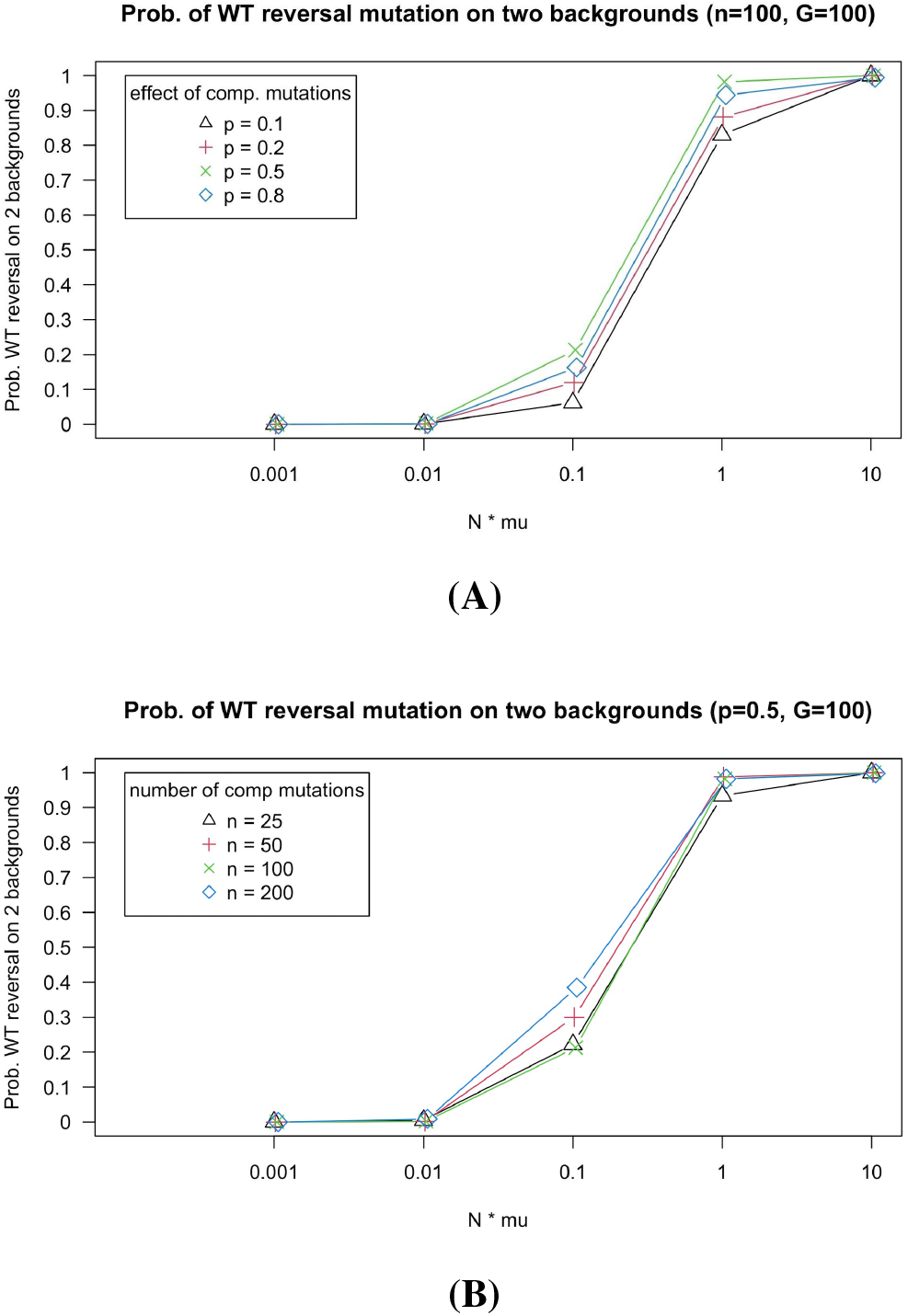
**(A)** The probability that the *WT* reversal mutation occurs on two different backgrounds, i.e., in a clone without and one with compensatory mutation. In these simulations, clones with compensatory mutations can acquire a second mutation (i.e., they can revert to *WT*). **(B)** Similar to A, but simulations are run for different values of n, the target size for compensatory mutations. Simulations are run for 100 generations. Other parameters: *N* = 10, 000, *c* = 0.15, *n* = 100.

### 3. What to expect after just 100 generations in a sample of size 10?

To get a better sense of what to expect in an experimental setting, we ran simulations for 100 generations and sampled 10 bacterial cells from the simulated populations. Figure 4A-E depicts the outcomes of 10 independent simulation runs for each of five different values of *N*µ*. We find very different outcomes, depending on the *N*µ* value (mutational supply in the population). Outcomes vary between replicate runs as well. Still, it is clear that some outcomes are more common for some values of *N*µ*.

**Figure 4:**
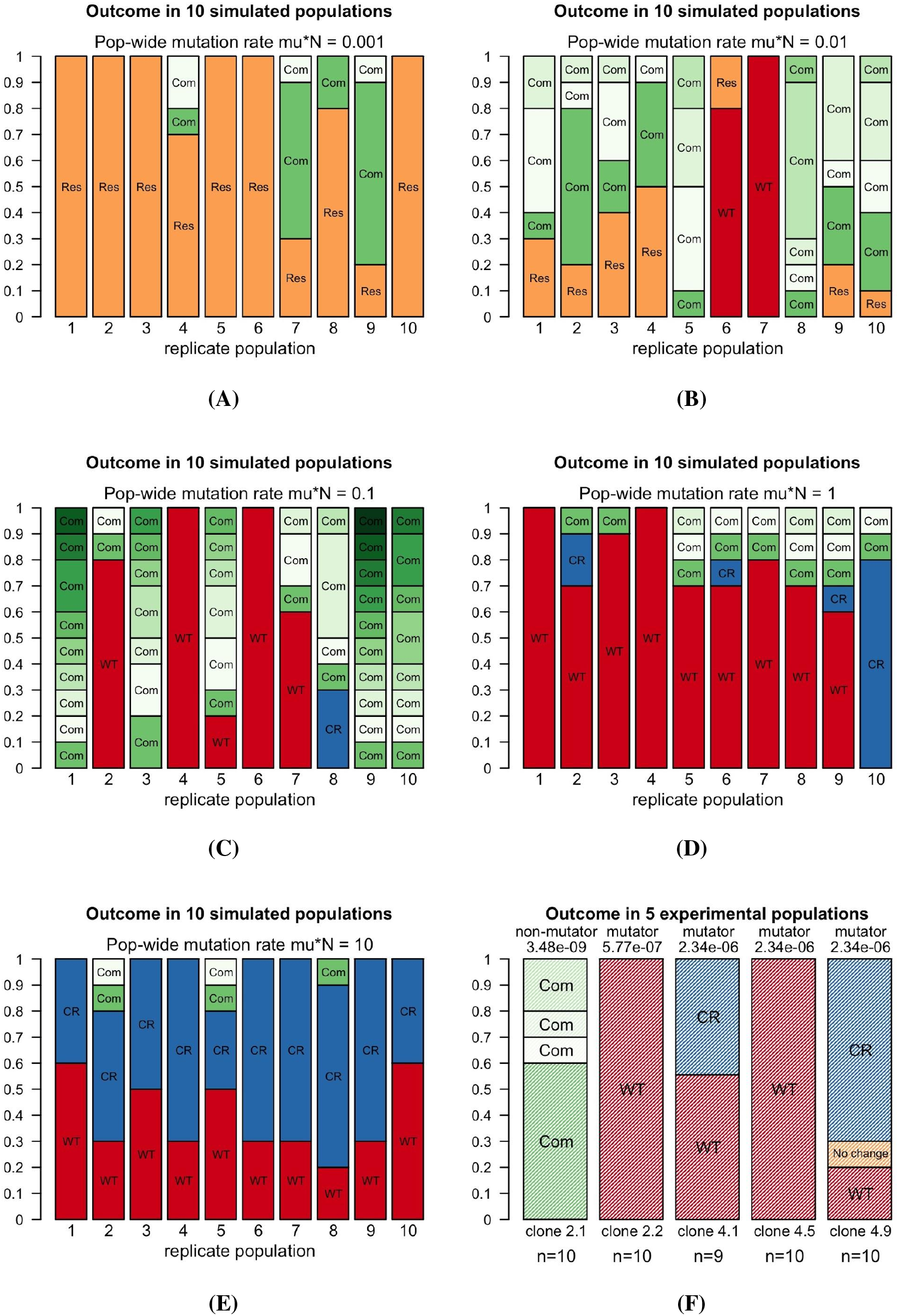
The dynamics of resistance, compensation and reversion in computational experiments A-E and previously published experimental results F. *Res* = resistant alleles; *Com* = compensated; *WT* = wild type; *CR* = compensated reversal. Different shades of green correspond to different compensatory mutations. Note that in 4F (the experimental results), the non-mutator yielded four types of compensatory mutations and no reversions, while the mutators yielded only reversions to the wild type, or combinations of reversions and compensatory mutations within the same population. There was one sample from clone 4.9 that didn’t have a reversal nor a compensatory mutation (indicated in orange). These results from the mutator clones are similar to what is observed in the computational experiments, when *N*µ* was set to 1 or 10, in Figure 4D-E. Other parameters: *N* = 10, 000, *c* = 0.15, *p* = 0.5, *n* = 100.

For these simulations all possible compensatory mutations have the same effect size and compensate half of the cost of the resistance mutation (comp = 0.5), so that the relative fitness values are 0.85 (for the resistant genotype), 0.925 (for the compensated genotype) and 1.0 (for the reverted WT genotype).

When *N*µ* equals 0.001 (Figure 4A), the populations do not evolve much over 100 generations and most of the sampled bacteria still carry the costly resistance mutation (indicated by the color orange and the word “Res”), with some compensatory mutations in some of the populations too (green colors).

When *N*µ* equals 0.01 (Figure 4B), it is more likely that compensatory mutations reach high frequencies (green colors). In a few cases, the *WT* reversal mutation reaches a high frequency too (red).

When *N*µ* equals 0.1 (Figure 4C) we observe reversal to *WT* (indicated by the color red and the word “*WT* ”) and compensatory mutations as well as a few bacteria with neither a reversal nor a compensatory mutation. We also see reversion combined with compensation (indicated by the color blue and the word “*CR*”), meaning that a compensatory mutation occurred prior to the reversion mutation that ultimately led to the loss of the costly adaptation.

When *N*µ* is 1 (Figure 4D), reversion is observed more frequently than compensation. When *N*µ* is 10 (Figure 4E), the most common outcome, by far, becomes reversion (in red and blue), leading to the loss of the costly adaptation.

In summary, we find that a 10 or 100-fold increase in mutation rates is sufficient to greatly modify probability of reversing the evolution of costly adaptive alleles. With lower mutation rates we see less reversion (with and without compensation) and with higher mutation rates we see more reversion (with and without compensation). When *N*µ* is 1 or 10, we see the WT reversal mutations on different backgrounds.

### 4. Comparison between computational results and previous experimental results

The results of our simulations correspond well with the experimental results that were previously observed in a study of costly adaptations in the RNA polymerase core enzyme in experimental *E. coli* long-term stationary phase (LTSP) populations. [23] In these experiments (see supplement), 5 clones with the (previously) adaptive mutation, which was shown to carry a cost to growth within fresh rich media, were used to initiate ∼ 100-generation serial dilution experiments, to see if they would lose the (previously) adaptive mutation or not. Four of the 5 clones were mutator clones that during LTSP had acquired a mutation within a mismatch repair gene, leading to substantially higher mutation rate. The fifth clone did not acquire such a mismatch repair mutation and thus had a normal mutation rate [29].

Results for the non-mutator clone (2.1) in the experiment (Figure 4F) correspond well to the simulations where *N*µ* = 0.01 or 0.1 (4B - 4C). In both simulations and experiments, several different compensatory mutations (in green) are observed in the population sample, but in most simulations and in the experiment no reversal is observed.

On the other hand, results for the four mutator clones were very different (Figure 4F). For three of the four mutator clones, all sequenced bacteria (9 or 10 per clone) had reverted to WT, whereas for one of the mutator clones (clone 4.9), 9 of 10 sequenced bacteria had reverted. This means that in most cases, the costly RNAPC adaptation was lost. These results are similar to what is observed in the computational experiments outlined in Figure 4D-E, when *N*µ* was set to 1 or 10.

The results of our simulations are consistent with previously published experimental results: we observe a shift from observing the persistence of costly adaptations through compensation, to the loss of costly adaptations due to reversion, for ∼ 100-fold changes in the rate of mutation.

### 5. Evolutionary dynamics change when fully compensating mutations are included

The simulations described above assumed that compensatory mutations do not fully alleviate costs associated with costly adaptations (*p <* 1). Next, we asked whether we would find similar results if one or a few of the compensatory mutations would fully restore fitness. We find that including perfectly compensating (*p* = 1) mutations in our simulations changes the outcomes drastically.

In figure 5 we show that when there are no perfectly compensating mutations, the fraction of the sample that carries the reversal mutation goes up with increasing mutation supply rate (*N*µ*). This can also be seen in Figure 4A-F. With higher mutation rates, we see more and more reversals.

However, when we include even just one fully compensating mutation, the average fraction of the sample that carries the reversal mutation is much lower. Specifically, when there is one perfectly compensating mutation (with the same mutation rate as the reversal mutation), on average half of the population will carry the compensatory mutation, the other half reversing to the wild type. When there are more than one perfectly compensating mutations, the expected fraction of the population that carries the reversal mutation decreases. As more perfectly compensating mutations are available, reversal to the wild type becomes less likely.

**Figure 5:**
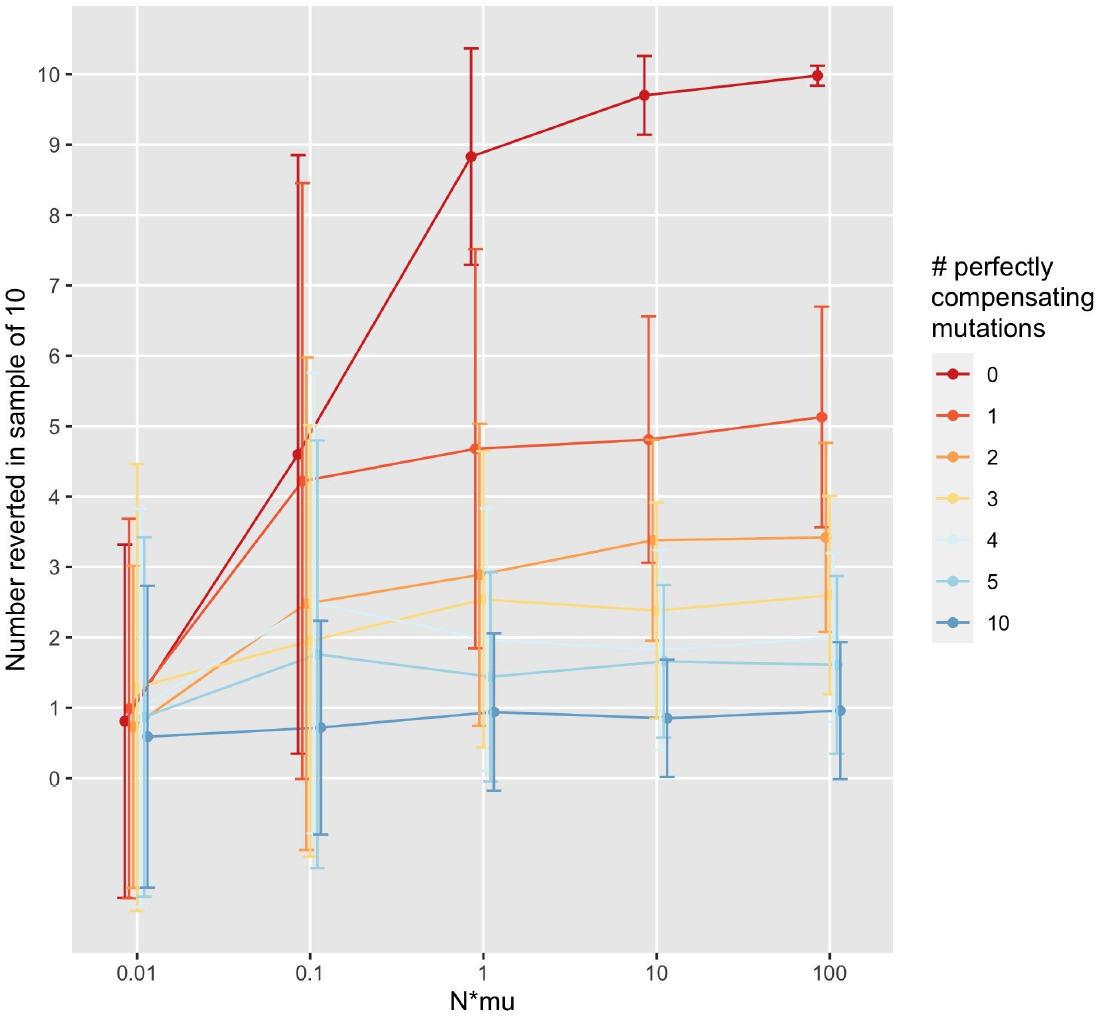
Simulations of evolution across mutation rates under different numbers of available fully-compensatory mutations. The x-axis corresponds to ranges of mutation supply (*N*µ*), and the y-axis shows the number of sampled bacteria where reversion to the wild type occurred. Here we observe that even the presence of a single fully compensatory mutation (dark orange line) greatly decreases the likelihood that the wild type allele will arise. The simulations were run for 100 generations. Other parameters: *N* = 10, 000, *c* = 0.15, *p* = 0.5, *n* = 100.

Note that in the experimental results (figure 4F), we observe that in the mutator populations, the reversion mutation is seen in nearly all sequenced bacteria, which is compatible with a model where none of the compensating mutations fully restore the fitness effects of resistance. This experimental outcome would be unlikely if there were any perfectly compensating mutations available.

In this manner, the simulation results outlined in figure 5 help to diagnose molecular evolutionary specifics of the empirical findings depicted in figure 4F. They suggest that in the experimental populations, there were no perfectly compensating mutations available. Even if only a single fully compensating mutation was available, we would have expected to see that only half (on average) of the population would have carried the reversal mutation and the other half would have carried a compensatory mutation only.

If compensatory mutations tend to only partially alleviate costs associated with adaptations, it is reasonable to expect that reversion mutations may ultimately occur and rise in frequency, even in clones that have already acquired a compensatory mutation. In our simulations of populations with higher mutation rates, we often observe clones that carry both a compensatory mutation and a reversion mutation (indicated in blue and with the letters “*CR*”). This suggests that reversion after partial compensation would also occur in lower-mutation-rate populations if we would follow the populations for (many) more generations.

## DISCUSSION

This study used population genetic theory and computational methods to examine the evolutionary dynamics of reversion and compensation in a context where mutations that confer adaptation to one setting confer a cost in another. Specifically, it sought to examine the conditions favouring reversion or compensation with regards to (1) differing mutation rates, (2) different values for the magnitude of fitness recovery for compensatory mutations.

Consistent with previously published laboratory findings, we find that several aspects of the evolutionary dynamics of resistance and compensation differ significantly at higher mutation rates [23]. At low mutation supply rates, the predominant genotypes are those that have remained resistant, and those that are resistant with standard compensatory mutations. In higher mutation rates settings, standard reversion (to the wild type) and compensated reversal (genotypes where both compensatory mutations and reversal mutations arise) are the predominant outcomes. It is important to note that in these settings, all resistance mutations have the same cost, and no compensatory mutations are fully restorative, which explains why reversion to the wild type can occur readily in high mutation rate contexts even after compensation. However, in settings where there are compensatory mutations that can fully restore fitness to wild type values (perfect compensation) in the original environment, reversion occurs less readily. This effect is especially visible in high mutation settings (see figure 5).

Our results are limited by the boundaries of our questions, and by extension, our modeling approach: we assume a simple, smooth fitness landscape, with no sign epistasis between mutations. In addition, resistance in our model is caused by a single mutation. This is important to consider because multistep reversion can be limited on rugged, complex fitness landscapes with sign epistasis between mutations [20], [21]. This study also focuses on the simple case of a single environmental shift, from one where a resistance mutation is beneficial, to one where it is detrimental. A growing literature examines situations reflective of a fitness “seascape,” where environments fluctuate, leading to capricious evolutionary dynamics [30], [31].

Nonetheless, our results highlight the utility in using theoretical and computational approaches to inform findings from experimental evolution of resistance, compensation, and reversion. They fortify notions that mutation rate is a strong modulator of evolutionary dynamics, a finding that is supported in studies of the clinical development of antibiotic resistance [29], [32]. Given this, the resistance management paradigm should continue to consider the effect varying mutation rates can have on the tendency of costly adaptations to persist or revert.

More broadly, future examinations of experimental evolution should carefully consider the mutation rate background in which these evolutionary dynamics occur, and more rigorously examine the population genetic (and mechanistic, if possible) underpinnings of compensation. These features can play an underappreciated role in the pace and direction of adaptive evolution, with implications that transcend the evolution of drug resistance or the microbial world.

## DATA AVAILABILITY

The experimental results discussed in the manuscript can be found in the original reference. [23] All code is available on github: https://github.com/pleunipennings/CompensatoryEvolution.

## ACKNOWLEDGMENTS

The authors would like to thank Joachim Krug and his group for discussion. The authors would also like to thank Victor A. Meszaros for editorial support.

