## Supplemental Appendix for "Reversion is most likely under high mutation supply, when compensatory mutations don’t fully restore fitness costs"

### Mathematical methods supplement

#### *Probability that the wild type arises before compensatory mutation fixes*

We calculate the probability that the $WT$ mutation occurs before fixation of the compensatory mutations using the following assumptions

- Bacteria can either acquire a compensatory mutation or a WT reversal mutation, not both.
- Absolute fitness of resistant bacteria $w_{res} = 1 - c$.
- Absolute fitness of the compensatory mutant $w_p = 1 - c(1 - p)$.
- We assume that compensatory mutations increase in frequency deterministically, starting from 1 individual, and therefore the number of resistant bacteria decreases deterministically, starting from N-1 following $N_{res}[t + 1] = N_{res}[t] \cdot w_{res}/\bar{w}[t] \cdot (1 - \mu)^n$ where $\bar{w}$ is the average population fitness.
- Now, for each generation, we have the number of resistant and compensated individuals and the average population fitness, which allows us to calculate selection coefficient and fixation probability for a WT reversal mutation.
- In a given generation, $t$, the selection coefficient of a WT reversal mutation is $s_{wt}[t] = w_{wt}/\bar{w}[t] - 1$ (where $w_{wt} = 1$) and the fixation probability of such mutation is $p_{fix}[t] = (1 - e^{(-2s_{wt}[t])})/(1 - e^{(-2Ns_{wt}[t])})$.
- The probability that at least one successful mutation occurs in a generation, $p_{wt}[t]$, is calculated as 1 minus the probability that none of the resistant individuals mutates and fixes. $p_{wt}[t] = 1 - (1 - \mu \cdot p_{fix}[t])^{N_{res}[t]}$
- The total probability of a successful WT reversal mutation before the compensatory mutations fix is calculated as a sum over all generations:
  $P_{tot} = \sum_{t=1}^{\infty}(p_{wt}[t]) \cdot \prod_{d=1}^{t-1}(1 - p_{wt}[d])$ where the product denotes the probability that no successful $WT$ mutation had occurred in previous generations.
- Even though we sum to infinity, the opportunity for $WT$ mutants in this model only lasts until the compensatory mutations have fixed. Over time, as the number of resistant (non-compensated) bacteria goes down, the mutational supply of $WT$ reversals goes down too (because here only resistant bacteria can mutate to WT). In addition, the population fitness ($\bar{w}$) goes up, so the selection coefficient ($s_{wt}$) and therefore the fixation probability $p_{fix}$ for $WT$ goes down.
- Note that our calculations somewhat overestimate the probability a successful $WT$ mutation occurs (see figure 3), likely because we overestimate $p_{fix}$. The reason may be that we assume that the fixation probability is determined by the selection coefficient in the generation that the mutation occurs. In reality, the selection coefficient in the following generations plays a role too, and since the population fitness is increasing, the selection coefficient is decreasing.

### EXPERIMENTAL SUPPLEMENT

For further clarity on the experimental results, we will briefly describe the results from 5 of the experimental E. coli populations as observed in [1]:

- Each of these 5 experimental populations was initiated from a clone with an adaptive mutation in the RNA polymerase core enzyme that increases fitness under long-term stationary phase – but that is costly in rich media. [1].
- One clone (2.1) had a normal mutation rate of $3.48 \cdot 10^{-9}$ per site. The other four clones were mutators, deficient in their mismatch repair, resulting in much higher mutation rates.
- Three of the clones (4.1, 4.5 and 4.9) had the same specific mutator allele, resulting in a mutation rate estimated to be 672-fold higher than that of the non-mutator clone.
- The remaining mutator clone (2.2). carried a different mutator allele resulting in a mutation rate 165-fold higher than that of the non-mutator.

These five experimental populations underwent daily dilutions into fresh media for 16 growth cycles, which corresponds to around 100 generations. During the serial dilution experiments, all five clones greatly improved the rate at which they grew exponentially in fresh media.

After 100 generations of evolution, 9-10 clones from each of the resulting populations were fully sequenced. This allowed us to examine whether the observed improvements in growth rate were achieved through compensation, reversion, or a combination of both. As can be seen, for the non-mutator populations, all 10 sequenced bacteria carried a compensatory mutation, while still maintaining their original costly mutation. While each clone carried a single compensatory mutation, four different compensatory mutations were observed within the population, indicating that this population adapted through a soft sweep [2]. These results were quite similar to what we observed in our simulations, when $N{*}\mu$ was set to 0.1 (Figure 4C) [3]. In the experiments with mutator strains, both compensatory and reversal mutations were observed (Figure 4F).
